# Coupled Co-clustering-based Unsupervised Transfer Learning for the Integrative Analysis of Single-Cell Genomic Data

**DOI:** 10.1101/2020.03.28.013938

**Authors:** Pengcheng Zeng, Jiaxuan WangWu, Zhixiang Lin

**Affiliations:** Department of Statistics, The Chinese University of Hong Kong, HK

**Keywords:** Clustering, Unsupervised transfer learning, Information-theoretic co-clustering, Single-cell genomic

## Abstract

Unsupervised methods, such as clustering methods, are essential to the analysis of single-cell genomic data. Most current clustering methods are designed for one data type only, such as scRNA-seq, scATAC-seq or sc-methylation data alone, and a few are developed for the integrative analysis of multiple data types. Integrative analysis of multimodal single-cell genomic data sets leverages the power in multiple data sets and can deepen the biological insight. We propose a coupled co-clustering-based unsupervised transfer learning algorithm (*couple*CoC) for the integrative analysis of multimodal single-cell data. Our proposed *couple*CoC builds upon the information theoretic co-clustering framework. We applied *couple*CoC for the integrative analysis of scATAC-seq and scRNA-seq data, sc-methylation and scRNA-seq data, and scRNA-seq data from mouse and human. We demonstrate that *couple*CoC improves the overall clustering performance and matches the cell subpopulations across multimodal single-cell genomic data sets. The software and data sets are available at https://github.com/cuhklinlab/coupleCoC.

## 1 Introduction

II Single-cell sequencing technologies have been developed to profile genomic features for many cells in parallel. Methods have been developed that can profile different types of genomic features, including single-cell RNA sequencing (scRNA-seq) that profiles transcription, single-cell ATAC sequencing (scATAC-seq) that identifies accessible chromatin regions, single-cell methylation assays that profile methylated regions. The available data sets are growing in the area of single-cell genomics (Rotem et al., 2015; Cusanovich et al., 2015; Rozenblatt-Rosen et al., 2017). Compared to bulk genomic data, single-cell genomic data is characterized by high technical variation and high noise level, due to the minimal amount of genomic materials isolated from individual cells (Kharchenko et al., 2014; Lun et al., 2016; Vallejos et al., 2017; Zhu et al., 2018; Hicks et al., 2018). These experimental factors make it challenging to analyze single-cell genomic data, and they can impact the result and interpretation of unsupervised learning methods, including dimension reduction and clustering (Jaitin et al., 2014; Usoskin et al., 2015; Lafon and Lee, 2006; Vandermaaten, 2008).

Most of the clustering methods developed for single-cell genomic data are focused on one data type. Here we first review the methods for clustering scRNA-seq, scATAC-seq or methylation data alone. DIMM-SC (Sun et al., 2017) builds upon a Dirichlet mixture model and aims to cluster droplet-based single cell transcriptomic data. RaceID (Grün et al., 2016) uses an iterative k-means clustering algorithm based on a similarity matrix of Pearson’s correlation coefficients. SIMLR (Wang et al., 2017) implements a kernel-based similarity learning algorithm, where RBF kernel was used with Euclidean distance. SC3 (Kiselev et al., 2017) and SAFE-clustering (Yang et al., 2018) belong to the class of ensemble clustering algorithms. The former combines the clustering outcomes of several other methods and the latter embeds several other methods and utilizes hypergraph-based partitioning algorithms. SOUP (Zhu et al., 2019) first identifies the set of pure cells by exploiting the block structures in cell-cell similarity matrix, uses them to build the membership matrix, and then estimates the soft memberships for the other cells. To cluster scATAC-seq data, *chrom*VAR (Schep et al., 2017) evaluates groups of peaks that share the same motifs or functional annotations together. *sc*ABC (Zamanighomi et al., 2018) weights cells by sequencing depth and applies weighted K-medoid clustering to reduce the impact of noisy cells with low sequencing depth. *cis*Topic (Gonzalez-Blas et al., 2019), applies latent Dirichlet allocation model on scATAC-seq data to identify *cis*-regulatory topics and simultaneously clusters cells and accessible regions based on the cell-topic and region-topic distributions. Xiong et al. (2019) proposed the SCALE method which combines a deep generative framework and a probabilistic Gaussian mixture model to learn latent features for scATAC-seq data. Clustering methods have also been proposed for methylation data (Shen et al., 2007; Siegmund et al., 2004). Siegmund et al. (2004) argued that model-based clustering techniques are often superior to nonparametric approaches. Houseman et al. (2008) proposed a model-based clustering method for methylation data produced by Illumina GoldenGate arrays. Their method incorporates a beta mixture model (Ji et al., 2005). A weighted model-based clustering called LmiWCluster for Illumina BeadArray was introduced in Kuan et al. (2010). It weights each observation according to the detection P-values systematically and avoids discarding subsets of the data. Kapourani and Sanguinetti (2019) proposed Melissa, a Bayesian hierarchical method to cluster cells based on local methylation patterns, and Melissa was shown to discover patterns of epigenetic variability between cells.

High technical variation and high noise level in single-cell genomic data sets motivate the development of integrative analysis methods, which combines multiple data sets. To identify sub-populations of cells that are present across multiple data sets, Butler et al. (2018) introduced an analytical strategy for integrating scRNA-seq data sets based on common sources of variation, enabling the identification of shared populations across data sets and downstream comparative analysis. In order to compare data sets across experiments, Kiselev et al. (2018) presented a method called SCMAP for projecting cells from an scRNA-seq data set onto cell types or individual cells from other experiments. Also, scVDMC (Zhang et al., 2018), a method of variance-driven multi-task clustering of single-cell RNA-seq data, was proposed to cluster single cells in multiple scRNA-seq experiments of similar cell types and markers but varying expression patterns, by utilizing multiple single-cell populations from biological replicates or different samples. Duren et al. (2018) developed *couple*NMF algorithm to address the increasingly common situation where two or more types of sc-genomics experiments are performed on different sub-samples from the same cell population. *couple*NMF is based on extensions of non-negative matrix factorization. The connection between chromatin accessibility and gene expression builds upon prediction models trained from bulk data with diverse cell types. Stuart et al. (2019) developed a strategy to “anchor” diverse data sets together with the capability of integrating single-cell measurements not only across scRNA-seq technologies, but also across different modalities. Their work presented a strategy for the assembly of harmonized references and transfer of information across data sets. Lin et al. (2019) proposed *sc*ACE, a model-based approach to jointly cluster single-cell chromatin accessibility and single-cell gene expression data. The model does not rely on training data to connect the two data types and allows for statistical inference of the cluster assignment. A more comprehensive discussion on integration of single-cell data across samples, experiments, and types of measurement is presented in David et al. (2020).

In this paper, we try to answer the following questions: can we cluster one type of single-cell genomic data (scATAC-seq data or single-cell methylation data) by incorporating the knowledge from another type (scRNA-seq data)? And can we cluster single-cell genomic data from one species (mouse scRNA-seq data) by learning patterns from another species (human scRNA-seq data)? These questions belong to the field of unsupervised transfer learning. Unsupervised transfer learning aims to improve the learning of one data set, called the target data, using the knowledge learnt in another data set, called the source data (Pan and Yang, 2009). Wang et al. (2008) proposed a transferred discriminative analysis (TDA) algorithm to solve the transfer dimensionality reduction problem. As a clustering version of the self-taught learning (Raina et al., 2007), Dai et al. (2008) proposed a self-taught learning algorithm (STC) to learn a common feature space across data sets, which improves clustering for the target data. Details on transfer learning are provided in the survey by Pan and Yang (2009).

Here we propose a novel coupled co-clustering-based (*couple*CoC) unsupervised transfer learning model for the integrative analysis of single-cell genomic data sets, as shown in Fig.1. Our goal is to use one dataset, the source data, to improve clustering of another dataset, the target data. We assume that features in the two datasets are linked: scRNA-seq and scATAC-seq data are connected by gene expression and promoter accessibility/gene activity score; scRNA-seq and single-cell methylation data are connected by gene expression and gene body hypomethylation; human and mouse scRNA-seq data are connected by homologs. We conduct co-clustering for the source data S and the target data T simultaneously (Fig.1(a)), where the linked features enable the transfer of knowledge between the source data and the target data. We also introduce a distribution-matching term of cell clusters that provides another level of knowledge sharing, and encourages the obtained clusters to be similar in source data and target data (Fig.1(b)). Last, the degree of knowledge transfer is learnt adaptively from the data.

**Figure 1:**
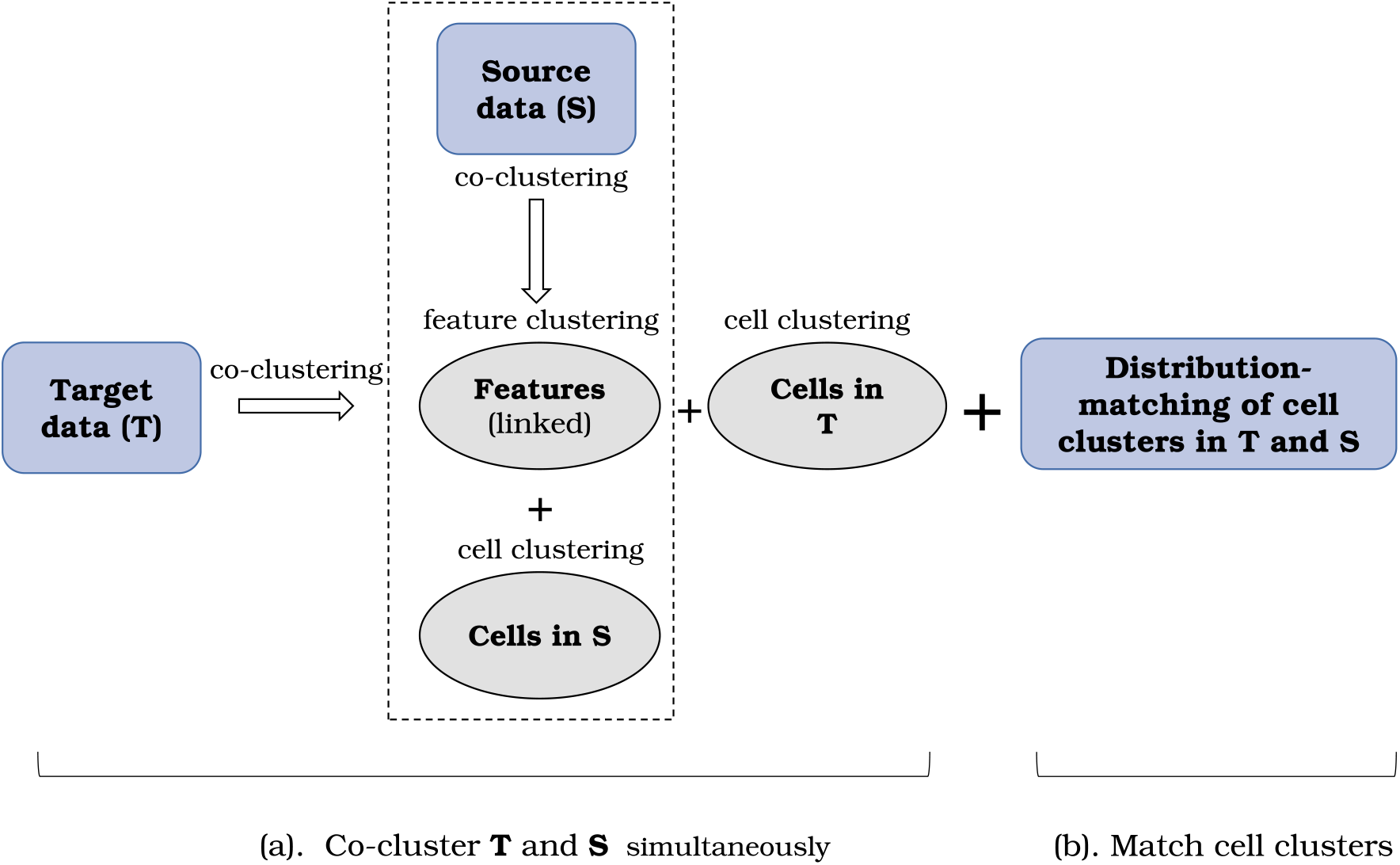
The schematic of the coupled co-clustering-based (*couple*CoC) unsupervised transfer learning model

The rest of the paper is organized as the following. We present the methods in Section 2. Simulation studies and real data analysis are presented in Section 3. The conclusion is presented in Section 4.

## 2 Methods

In this section, we first formulate the problem and then introduce the *couple*CoC algorithm, followed by approaches to selecting features, pre-processing the data, determining the number of cell clusters and tuning the parameters.

### 2.1 Problem Formulation

We first pre-process the data matrices of scRNA-seq, scATAC-seq and single-cell methylation (details presented in Section 2.3). Depending on the implementation and the strength of the signal in separating the cells, we choose one dataset as the source data and the other dataset as the target data. More details on choosing the source data and target data are presented later in the real data section. Our goal is to use source data to improve the clustering of target data. We denote the target data as a matrix 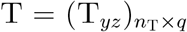, where T_*yz*_ represents the activity of the *z*-th feature in the *y*-th cell: the more active the feature, the higher the value. In scRNA-seq data, an active feature means the gene is expressed; in scATAC-seq data, an active feature means the promoter is accessible as accessible promoter is associated with active gene expression; in single-cell methylation data, an active feature means the gene body is hypomethylated as hypomethylated gene body is associated with active gene expression. Similarly, we denote the source data as a matrix 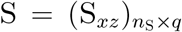, and the definition of an active feature is the same as that in target data. In this work, we use promoter accessibility to connect scATAC-seq data to scRNA-seq data.

The promoter accessibility can be easily replaced by gene activity score (Hannah et al., 2018) without changing the *couple*CoC algorithm. The gene activity score summarizes the accessibility of the regions near the gene body. The positive relation between promoter accessibility and gene expression is shown in Fig.5–7. The negative relation between gene body methylation and gene expression is shown in Fig.8. The positive relation between human and mouse homologs is shown in Fig.9.

We first use a toy example (Fig.2(a)) to illustrate our proposed method. We regard two different 4 × 6 matrices as the target data (T) and the source data (S), respectively. Let *Y* be a discrete random variable, which takes value from the set of cell indexes {1,…, *n*_T_} in target data T. Let *X* be a discrete random variable, which takes value from the set of cell indexes {1,…, *n*_S_} in source data S. Further, let *Z* be a discrete random variable, which takes value from the set of feature indexes {1,…, *q*}. Take the target data (Fig.2(a)) as an example: *Y* = *y*, where *y* = 1, … , 4, means that the selected cell is the *y*-th cell among 4 cells, and *Z* = *z*, where *z* = 1, …, 6, means that the selected feature is the *z*-th feature among 6 features. Let *p*_T_(*Y, Z*) be the joint probability distribution for *Y* and *Z*, which can be represented by a *n*_T_ × *q* matrix. *p*_T_(*Y* = *y, Z* = *z*) is the probability of the *z*-th feature being active in the *y*-th cell, and it is estimated from the observed target data T:

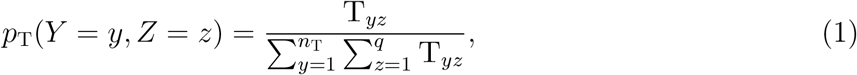

where *y* ∈ {1, …, *n*_T_}, *z* ∈ {1, …, *q*}. The marginal probability distributions are then expressed as 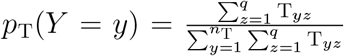, and 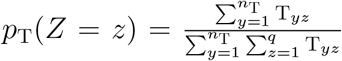. For the source data, *p*_S_(*X, Z*) is the joint probability distribution for *X* and *Z*; *p*_S_(*X*) and *p*_S_(*Z*) are the marginal probabilities. The distributions *p*_S_(*X, Z*), *p*_S_(*X*) and *p*_S_(*Z*) are calculated similarly to that in the target data.

**Figure 2:**
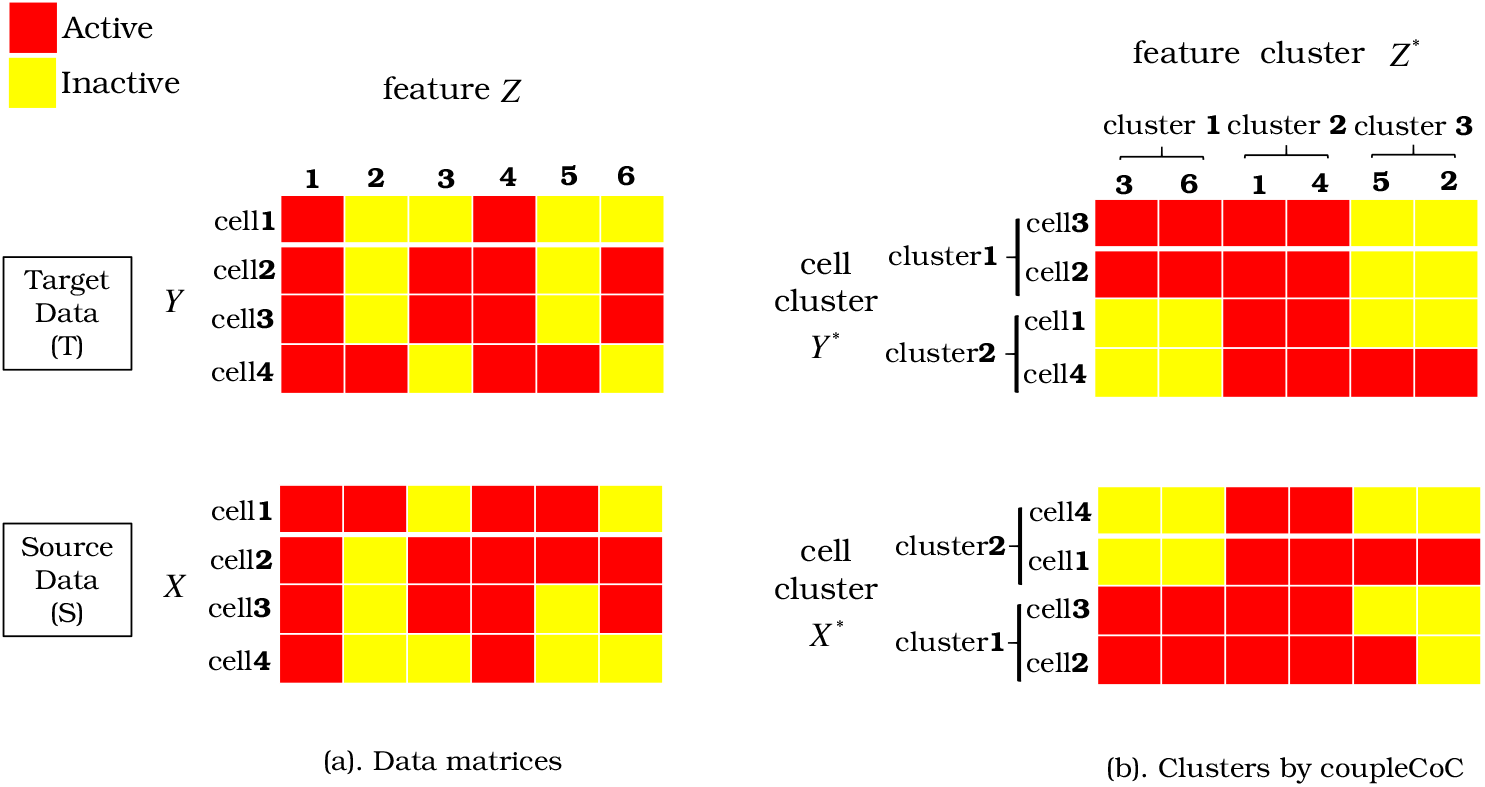
Toy example of our method (coupleCoC) for the analysis of single-cell data. (a). Red color means the corresponding features are active, and yellow color means inactive. (b). Data after clustering: cells and features are reordered such that similar cells and similar features are grouped together. Note that the clusters of cells are matched in the target data and source data by our method.

We aim at grouping similar cells into clusters and at grouping similar features into clusters as well. Suppose we want to cluster the cells in target data into *N*_T_ clusters, the cells in source data into *N*_S_ clusters, and the linked features into *K* clusters. Note that the values of *N*_T_ and *N*_S_ are pre-determined and *N*_T_ may not be equal to *N*_S_. *K* is a hyper-parameter. How the values of *N*_T_, *N*_S_ and *K* are determined will be presented in Section 2.4. Let the random variables *Y*\* and *X*\* take values from the set of cell cluster indexes {1, …, *N*_T_} and {1, …, *N*_S_}, respectively. We use *C*_*Y*_(·) to represent the clustering function for target data and *C*_*Y*_(*y*) = *i* (*i* = 1, …, *N*_T_) indicates that cell *y* belongs to cluster *i*. For source data, the clustering function *C*_*X*_(·) is defined similarly as that for the target data. Let the random variable *Z*\* take values from the set of feature cluster indexes {1, …, *K*}. We use *C*_*Z*_(·) to represent the clustering function of the features and *C*_*Z*_(*z*) = *j* (*j* = 1, …, *K*) indicates that feature *z* belongs to cluster *j*. The tuples (*C*_*Y*_(*y*), *C*_*Z*_(*z*)) and (*C*_*X*_(*x*), *C*_*Z*_(*z*)) are referred as co-clustering (Dhillon et al., 2003).

Let *p*_T_(*Y*\*, *Z*\*) be the joint probability distribution for *Y*\* and *Z*\*, which can be represented by a *N*_T_ × *K* matrix. Based on the Equation (1), this distribution can be expressed as

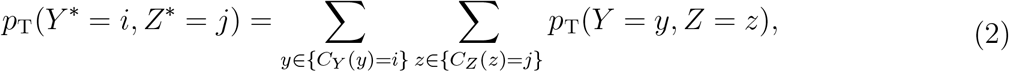

The marginal probability distributions are then expressed as 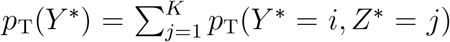 and 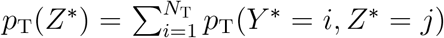. For the source data, *p*_S_(*X*\*, *Z*\*), *p*_S_(*X*\*) and *p*_S_(*Z*\*) are defined and calculated analogously to those for the target data.

The goal of coupled co-clustering-based unsupervised transfer learning in this work is to find the optimal cell clustering function *C*_*Y*_ for the cells in target data by utilizing the information of both the cell clustering function *C*_*X*_ for the cells in source data and the feature clustering function *C*_*Z*_.

### 2.2 The CoupleCoC Algorithm

Our proposed method extends the information theoretic co-clustering framework (Dhillon et al., 2003), which minimizes the loss in mutual information after co-clustering of instances and features. For target data T and source data S, the loss of mutual information after co-clustering can be expressed as

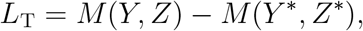

and

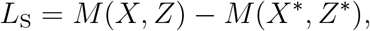

respectively, where *M*(·, ·) denotes the mutual information between two random variables: 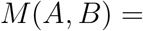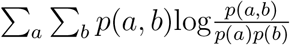. In practice, we only need to consider the elements satisfying *p*(*a, b*) ≠ 0.

We propose the following objective function for coupled co-clustering of the target data T and the source data S:

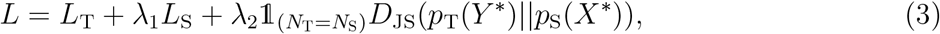

where 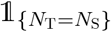 equals to 1 if *N*_T_ = *N*_S_ and 0 otherwise, and *D*_JS_(*p*_T_(*Y*\*)‖*p*_S_(*X*\*)) is the Jensen-Shannon Divergence, defined as

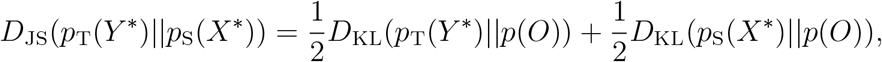

where 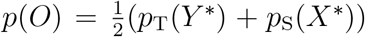, and *D*_KL_(·‖·) denotes the Kullback-Leibler divergence (KL divergence) between two probability distributions (Cover and Thomas, 1991), where 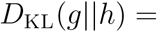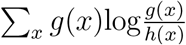. The term *L*_T_ is specific to the target data T and the term *L*_S_ is specific to the source data S. The two terms *L*_T_ and *L*_S_ share the same feature clusters *C*_*Z*_, and *Z*\* can be viewed as a bridge to transfer knowledge between T and S. λ_1_ is a hyperparameter that controls the contribution of the source data S to the target data T. Another component in Equation (3) that bridges T and S is the distribution-matching term *D*_JS_(*p*_T_(*Y*\*)‖*p*_S_(*X*\*)). The term *D*_JS_(*p*_T_(*Y*\*)‖*p*_S_(*X*\*)) encourages *Y* * and *X*\*, the clusters in target and source data, to be similar. On top of improving the clustering accuracy, this term can offer better matching of the clusters in target and source data. When *N*_T_ ≠ *N*_S_, the term *D*_JS_(*p*_T_(*Y*\*) *p*_S_(*X*\*)) disappears, and the matching of cell types can be carried out by post-processing the cluster labels, i.e., swapping cluster labels for each data type and choosing subsets of clusters that minimize 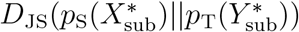, where the number of cluster labels in 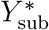 is equal to that in 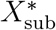 (an example is presented in Section 3.2.3). λ_2_ is a hyperparameter on how well the clusters are matched in target and source data. The tuning of hyperparameters λ_1_ and λ_2_ will be presented in Section 2.4. We call our proposed method *couple*CoC, which is the abbreviation of coupled co-clustering.

We need to solve the optimization problem:

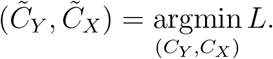

This optimization problem is non-convex and challenging to optimize. We can rewrite the terms *L*_T_ and *L*_S_ in the objective function *L* into the forms of KL divergence and the reformulated objective function is easier to optimize. More specifically, for the target data T,

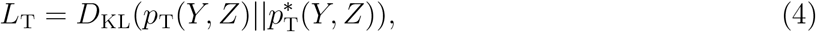

where 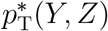 is defined as

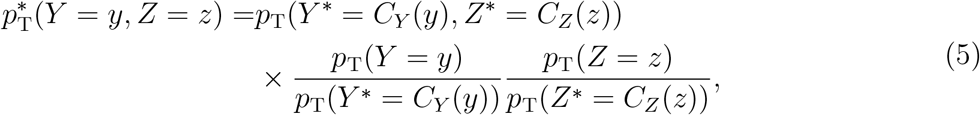

The probabilities *p*_T_(*Y*\* = *C*_*Y*_(*y*), *Z*\* = *C*_*Z*_(*z*)), *p*_T_(*Y* = *y*), *p*_T_(*Y*\* = *C*_*Y*_ (*y*)), *p*_T_(*Z* = *z*), *p*_T_(*Z*\* = *C*_*Z*_(*z*)) are estimated as in the previous sections. Further, we have

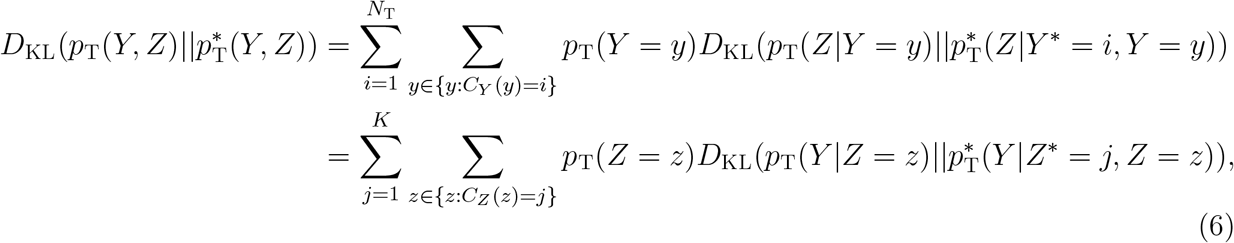

where 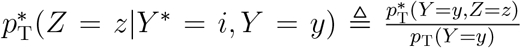, *z* = 1, …, *q*, for any *y* ∈ {*y* : *C*_*Y*_(*y*) = *i*}, and 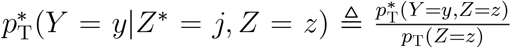, *y* = 1, …, *n*_T_, for any *z* ∈ {*z* : *C*_*Z*_(*z*) = *j*}. Details on the proof of Equations (4) and (6) are presented in Dhillon et al. (2003) and Dai et al. (2008). For the source data S, we can rewrite *L*_S_ similarly as we rewrote *L*_T_ in Equations (4), (5) and (6). Therefore, the objective function in Equation (3) can be rewritten as:

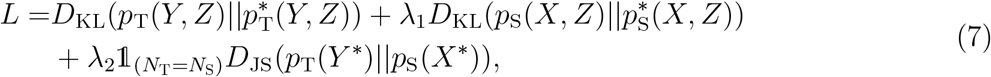

To minimize *L*, we can iteratively update the clustering functions *C*_*Y*_, *C*_*X*_ and *C*_*Z*_ as follows:

- Given *C*_*X*_ and *C*_*Z*_, update *C*_*Y*_. Minimizing *L* is equivalent to minimizing

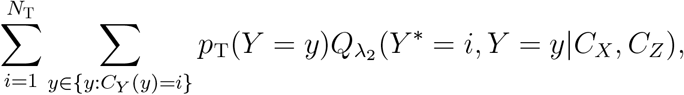

where

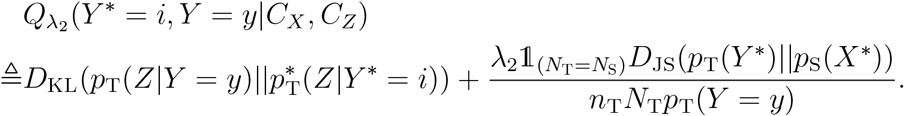 We iteratively update the cluster assignment *C*_*Y*_(*y*) for each cell *y* (*y* = 1, …, *n*_T_) in the target data, fixing the cluster assignment for the other cells:

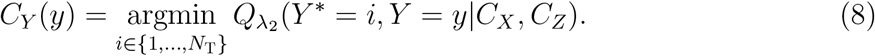
- Given *C*_*Y*_ and *C*_*Z*_, update *C*_*X*_. Minimizing *L* is equivalent to minimizing

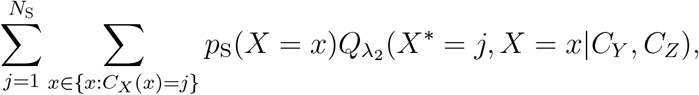

where

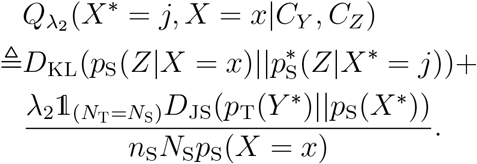 We iteratively update the cluster assignment *C*_*X*_(*x*) for each cell *x* (*x* = 1, …, *n*_S_) in the source data, fixing the cluster assignment for the other cells:

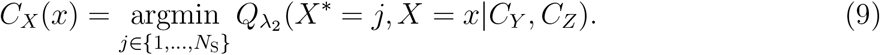
- Given *C*_*Y*_ and *C*_*X*_, update *C*_*Z*_. Minimizing *L* is equivalent to minimizing

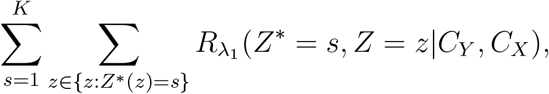

where

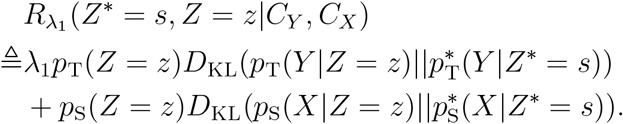 We iteratively update the cluster assignment *C*_*Z*_(*z*) for each feature *z* (*z* = 1, …, *q*), fixing the cluster assignment for the other features:

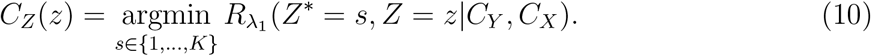

To summarize, the procedures of optimizing the global objective function (3) are as follows:

(I) Initialization. Calculate *p*_T_(*Y, Z*) and *p*_S_(*X, Z*) using the target data T and source data S. Initialize the clustering functions 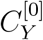, 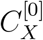 and 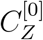. Initialize 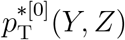 and 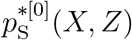.
(I) Iterate steps a and b until convergence.

a. Fix 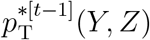 and 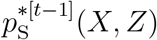 and sequentially update 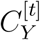, 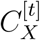 and 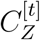 based on the Equations (8), (9) and (10).
b. Fix 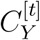, 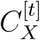 and 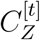 and update 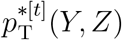 and 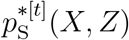.
(I) Output the clustering results *C*_*Y*_, *C*_*X*_ and *C*_*Z*_ in the last iteration.

The objective function (Equation (3)) is non-increasing in the updates of *C*_*Y*_, *C*_*X*_ and *C*_*Z*_, and the algorithm will converge to a local minimum. Finding the global optimal solution is NP-hard. Additionally, the *couple*CoC algorithm converges in a finite number of iterations due to the finite search space. In practice, this algorithm works well in both simulations and real data analysis.

### 2.3 Feature selection and data preprocessing

We first select the linked features: we use promoter accessibility for scATAC-seq data, gene body methylation for sc-methylation data, homologs for the human and mouse scRNA-seq data. We implement feature selection on the linked features before performing clustering. This pre-processing step speeds up computation and reduces the noise level. We used the R toolkit Seurat (Butler et al., 2018; Stuart et al., 2019) to select 1000 most variable features for each dataset.

After feature selection, we used log2(TPM+1) as the input for the scRNA-seq data, raw read count as the input for the scATAC-seq data and gene body methylation level as the input for the sc-methylation data. We imputed the missing values in sc-methylation data with the overall mean. We then standardized the target data T by minimax normalization:

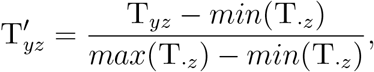

where *max*(T_·*z*_) and *min*(T_·*z*_) are the maximal and minimal expression level (or read counts) for feature *z* across all cells in the target data. We implemented the same minimax normalization for the source data S. Since the relationship between gene body methylation and gene expression is negative (Welch et al., 2019), we further transform the normalized sc-methylation data by 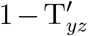.

### 2.4 Determining the number of cell clusters and parameter tuning

We use the Calinski-Harabasz index (Calinski and Harabasz, 1974) to pre-determine the number of cell clusters *N*_T_ for the target data and the number of cell clusters *N*_S_ for the source data separately, before we implement the *couple*CoC algorithm. Calinski-Harabasz index is proportional to the ratio of the between-clusters dispersion and the within-cluster dispersion:

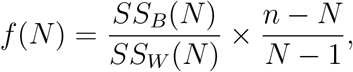

where *SS*_*B*_(*N*) is the overall between-cluster variance, and *SS*_*W*_(*N*) is the overall within-cluster variance, *N* is the number of cell clusters, and *n* is the total number of cells. For each cluster number *N*, we first cluster the dataset by minimizing the objective function *L*_T_ for the target data or *L*_S_ for the source data using the *couple*CoC algorithm where we set λ_1_ = λ_2_ = 0, and we then calculate *SS*_*B*_(*N*), *SS*_*W*_(*N*), and obtain *f*(*N*). We choose the number of cell clusters *N* with the highest Calinski-Harabasz index.

Our algorithm *couple*CoC has three tuning parameters: λ_1_, λ_2_ and K, which is the number of clusters for the features. It is hard to determine these paramters theoretically, and they are tuned empirically by grid search in practice (Dai et al., 2008). We use grid search to choose the best combination of hyperparameters that has the the highest Calinski-Harabasz index for the target data:

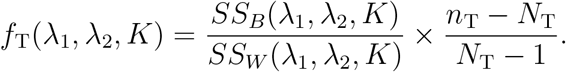

Here we only evaluate the clustering performance of the target data, because in *couple*CoC we expect to improve the clustering performance of the target data by learning knowledge from the source data. For each combination of hyperparameters 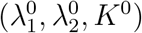, we first cluster the coupled datasets by minimizing the objective function *L*, and we then calculate 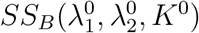, 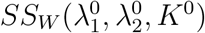, and obtain 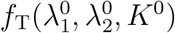, for the target data. We choose the search domains λ_1_ ∈ [0, 5], λ_2_ ∈ [0, 5] and *K* ∈ (0, 10). Grid search performs well in both simulated data and real data.

## 3 Results

### 3.1 Simulation studies

We follow the simulation setup given in Lin et al. (2019). The steps of generating T and S are given in Supplementary Text. We set the number of cells in both target data T and source data S as *n*_T_ = *n*_S_ = 100, and set the number of features as *q* = 100. We varied the differential degree (*w*) across clusters and the standard deviation (*σ*) of the generative distribution. Larger *w* leads to better separation of the cell clusters, and larger *σ* leads to higher noise level. We consider three different simulation settings, varying the parameters *w* and *σ*.

Table 1 presents the simulation results. We use four criteria to quantify the clustering results: purity, Rand index (RI), adjusted Rand index (ARI) and normalized mutual information (NMI) (the details of these criterion are given in Christopher et al. (2008)). The tuning parameters are set as λ_1_ = λ_2_ = 1.5, *K* = 3 by grid search. The trends for different clustering criteria are similar over the three settings. Compared with the first setting (*w* = 0.8, *σ* = 0.6), the second setting has a larger variance (*σ* = 1.2) in the target data, and the third setting has lower differential ability across the cell clusters (*w* = 0.7). Settings 2 and 3 have lower overall clustering performance compared with that in setting 1. As expected, our model (*couple*CoC) achieves better clustering results, compared with *k*-means, spectral clustering and hierarchial clustering. This is likely because of the fact that our model transfers the knowledge from the source data, which helps to improve the clustering performance on the target data. The other three methods do not utilize the information from the source data.

**Table 1:** The simulation results of clustering scRNA-seq data for 50 independent runs are summarized.

| Settings | Methods | Purity | RI | ARI | NMI |
| --- | --- | --- | --- | --- | --- |
| Setting 1<br>$w = 0.8, \sigma = 0.6$ | *coupleCoC* | 0.891 | 0.806 | 0.612 | 0.531 |
| | $k$-means | 0.811 | 0.706 | 0.411 | 0.352 |
|  | spectral clustering | 0.548 | 0.501 | 0.001 | 0.037 |
|  | hierarchical clustering | 0.549 | 0.501 | 0.001 | 0.035 |
| Setting 2<br>$w = 0.8, \sigma = 1.2$ | *coupleCoC* | 0.838 | 0.729 | 0.459 | 0.381 |
| | $k$-means | 0.701 | 0.594 | 0.188 | 0.162 |
|  | spectral clustering | 0.555 | 0.502 | -0.000 | 0.036 |
|  | hierarchical clustering | 0.557 | 0.503 | 0.002 | 0.040 |
| Setting 3<br>$w = 0.7, \sigma = 0.6$ | *coupleCoC* | 0.693 | 0.585 | 0.169 | 0.137 |
| | $k$-means | 0.620 | 0.533 | 0.067 | 0.061 |
|  | spectral clustering | 0.548 | 0.501 | 0.000 | 0.036 |
|  | hierarchical clustering | 0.550 | 0.501 | 0.001 | 0.037 |

### 3.2 Application to real data

In this section, we will present three real data examples, including one example with three cases for clustering human scATAC-seq data and scRNA-seq data, one example for clustering mouse sc-methylation data and scRNA-seq data, and one example with two cases for clustering human and mouse scRNA-seq data (drop-seq platform). Table 2 shows the details of the datasets included. In *couple*CoC, the dataset that is less noisy should be chosen as the source data. In examples 1 and 2, scRNA-seq data tends to be less noisy compared with scATAC-seq and sc-methylation data, so we chose scRNA-seq data as the source data. In example 3, the human data tends to be less sparse compared with the mouse data, so we chose the human data as the source data. We compare our method *couple*CoC with *k*-means, and commonly used clustering methods for single-cell genomic data, including SC3 (Kiselev et al., 2017), SIMLR (Wang et al., 2017), SOUP (Zhu et al., 2019) and Seurat (Stuart et al., 2019). We determined the number of cell clusters for *couple*CoC by the Calinski-Harabasz index, and we used the true number of cell clusters for the other methods except for the method Seurat, which automatically determines the number of cell clusters. Other than the clustering table, we use purity, Rand index (RI), adjusted Rand index (ARI) and normalized mutual information (NMI) to evaluate the clustering results.

**Table 2:** Datasets used in the real data analysis.

| Example No. | data types | # of cell types | # of cells | % of zero entry | data source |
| --- | --- | --- | --- | --- | --- |
| Example 1 (A) | scATAC-seq data (T) | 2 | 314 | 83.74% | Buenrostro et al. (2015) |
|  | scRNA-seq data (S) | 2 | 96 | 64.31% | Pollen et al. (2014) |
| Example 1 (B) | scATAC-seq data (T) | 2 | 1039 | 88.06% | Buenrostro et al. (2015) (GSE65360) |
|  | scRNA-seq data (S) | 2 | 201 | 57.36% | Buenrostro et al. (2015) (GSE65360) |
| Example 1 (C) | scATAC-seq data (T) | 3 | 1353 | 86.84% | Buenrostro et al. (2015) (GSE65360) |
|  | scRNA-seq data (S) | 3 | 297 | 58.64% | Buenrostro et al. (2015) (GSE65360); Pollen et al. (2014) |
| Example 2 | sc-methylation data (T) | 2 | 1102 | 21.80%(missing data) | Welch et al. (2019) (GSE126836) |
|  | scRNA-seq data (S) | 2 | 2383 | 82.86% | Welch et al. (2019) (GSE126836) |
| Example 3 (A) | mouse scRNA-seq data (T) | 2 | 113 | 92.50% | Fran et al. (2019) (*panglaodb.se*, SRS4237518) |
|  | human scRNA-seq data (S) | 2 | 171 | 88.90% | Fran et al. (2019) (*panglaodb.se*, SRS4660846) |
| Example 3 (B) | mouse scRNA-seq data (T) | 3 | 292 | 95.00% | Fran et al. (2019) (*panglaodb.se*, SRS4237518) |
|  | human scRNA-seq data (S) | 2 | 171 | 88.90% | Fran et al. (2019) (*panglaodb.se*, SRS4660846) |

#### 3.2.1 Example 1: clustering K562, HL60 and GM12878 cells of human scATAC-seq and scRNA-seq data

In this example, we evaluate our *couple*CoC method by clustering cell types K562, HL60 and GM12878 of human scRNA-seq and scATAC-seq data (Buenrostro et al., 2015; Pollen et al., 2014). We consider three cases: (A) 223 K562 and 91 HL60 scATAC-seq cells and 42 K562 and 54 HL60 deeply sequenced scRNA-seq cells; (B) 373 GM12878 and 666 K562 scATAC-seq cells, and 128 GM12878 cells and 73 K562 scRNA-seq cells; (C) all the cells in cases A and B. Note that the scRNA-seq datasets in cases A and B come from two different studies. The true cell labels are used as a benchmark for evaluating the performance of the clustering methods. Fig.10 shows the optimal number of cell clusters are *N*_T_ = 2 and *N*_S_ = 2 for both case A and case B and *N*_T_ = 3 and *N*_S_ = 3 for case C. The tuning parameters in *couple*CoC are set as λ_1_ = 1.5, λ_2_ = 2, *K* = 5 for case A, λ_1_ = 1, λ_2_ = 1.2, *K* = 5 for case B, and λ_1_ = 1, λ_2_ = 1, *K* = 5 for case C by grid search. The clustering results (Table 3) show that all six methods perfectly separated the cell types in the source data (scRNA-seq data), except for *k*-means in case C. Compared with *couple*CoC, the three methods, *k*-means, SIMLR and Seurat do not cluster the target data (scATAC-seq data) well in all three cases, SC3 does not perform well in cases B and C, and SOUP does not perform well in case C. This may be due to the fact that *couple*CoC utilize information from scRNA-seq data to improve the clustering of scATAC-seq data. SC3, SIMLR, and SOUP are designed for clustering one data type, so they cannot match cell types across the source data and target data. Our proposed method, *couple*CoC, has good clustering performance and the cell types are correctly matched. The clustering result of case C is not as good as that of cases A and B, which may be due to the fact that GM12878 is similar to HL60. Visualization of the co-clustering results by *couple*CoC is presented in Fig.3(a)-(c), where we grouped and segmented cell clusters and feature clusters by blue lines. Our proposed *couple*CoC clearly clusters similar cells and features in cases A and B, but not as well in case C.

**Table 3:**
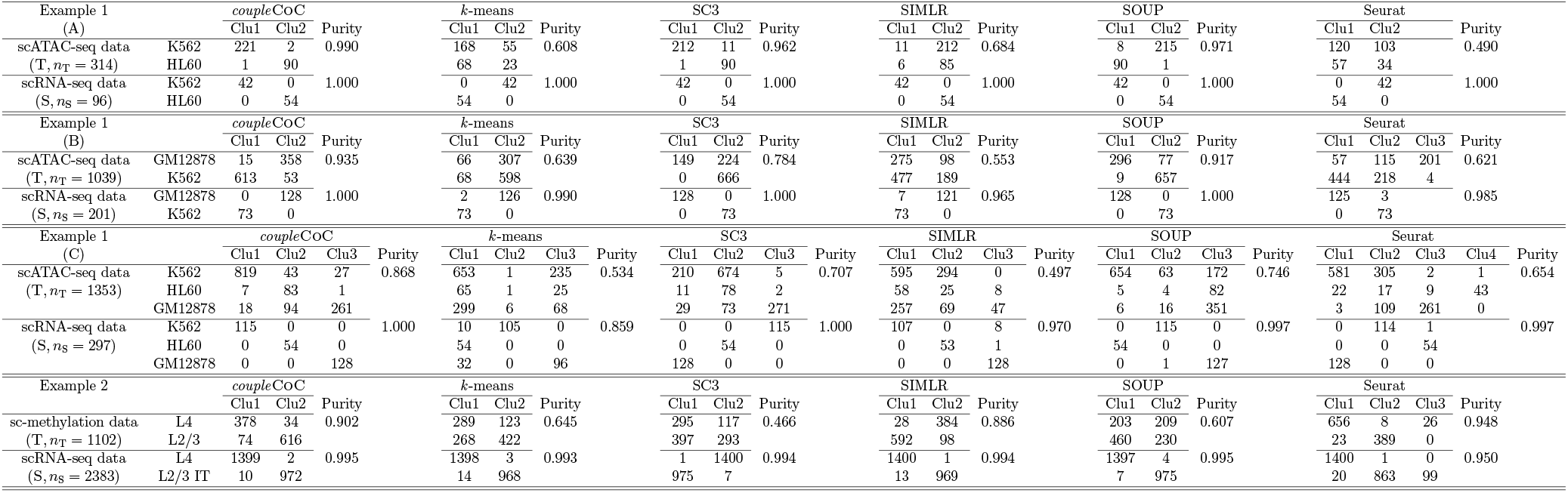
Clustering table and purity by *couple*CoC, *k*-means, SC3, SIMLR, SOUP and Seurat on both target data T and source data S for examples 1 and 2. “Clu” is the abbreviation of “Cluster”.

**Figure 3:**
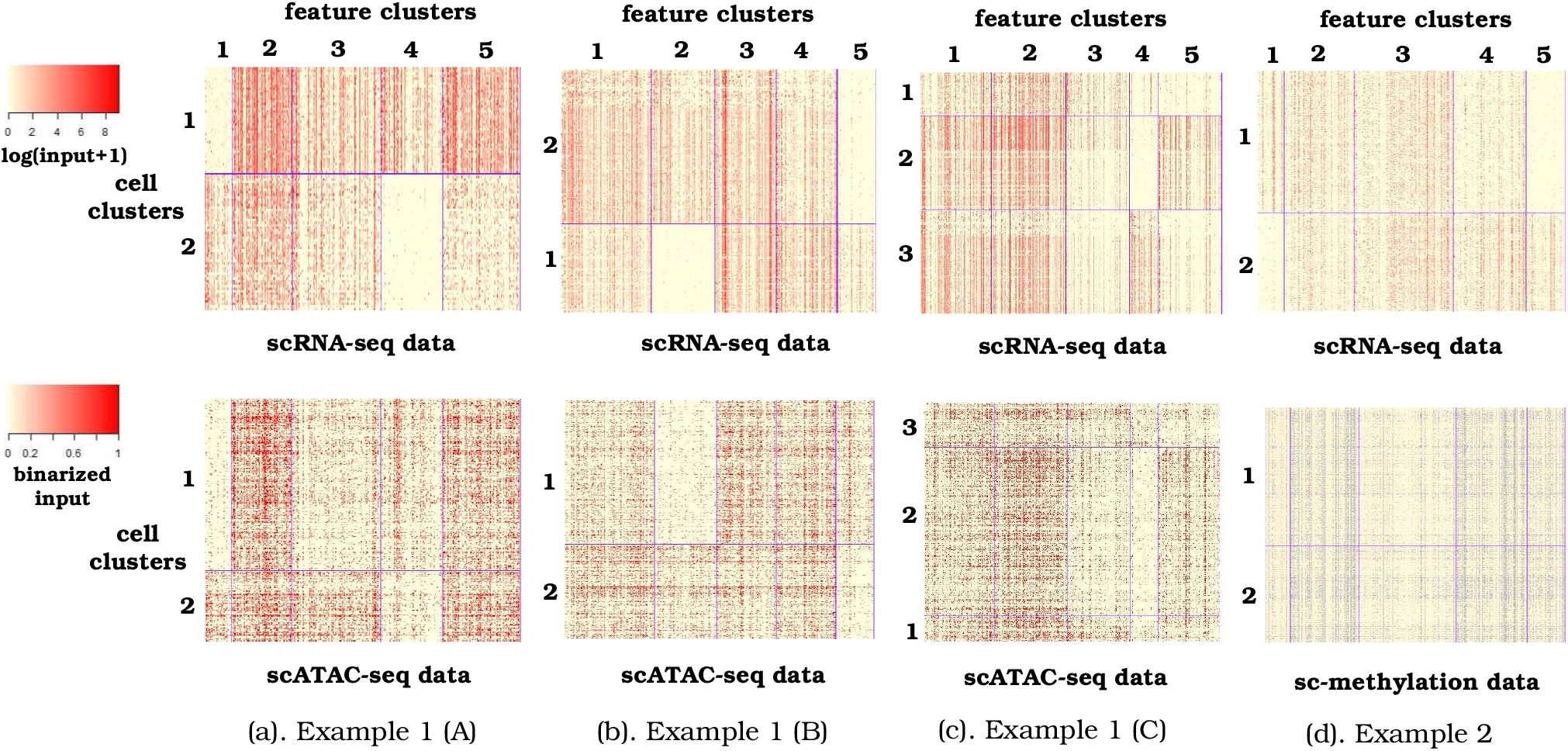
Heatmaps of the clustering results by *couple*CoC for examples 1 and 2. The input for scRNA-seq data are TPM. The input for scATAC-seq data and sc-methylation data are the raw read count and gene body methylation level, respectively, and both are binarized for visualization purpose. Grey color in the heatmap of sc-methylation data shown in (d) corresponds to missing data.

#### 3.2.2 Example 2: clustering frontal cortex L4 and L2/3 cells of mouse sc-methylation and scRNA-seq data

In the second example, we evaluate our method *couple*CoC by jointly clustering mouse frontal cortex scRNA-seq sc-methylation data and scRNA-seq data (Welch et al., 2019). We collected 412 L4 and 690 L2/3 sc-methylation cells, and 1401 L4 and 982 L2/3 IT scRNA-seq cells (“L4” and “L2/3” stand for different neocortical layers; IT is the abbreviation of intratelencephalic neuron.). The provided cell type labels are used as a benchmark for evaluating the performance of the clustering methods. Fig.10 shows the optimal number of cell clusters are *N*_T_ = 2 and *N*_S_ = 2. The tuning parameters in *couple*CoC are set as λ_1_ = 1, λ_2_ = 3, *K* = 5 by grid search. Example 2 in Table 3 shows that all six methods except Seurat have good clustering performance for scRNA-seq data. Second, our method *couple*CoC and SIMLR perform much better than the other methods for clustering sc-methylation data, and *couple*CoC matches well the cell types across the two data types. Fig.3(d) is the corresponding heatmap after clustering by *couple*CoC. For the scRNA-seq data, our proposed *couple*CoC clearly clusters similar cells and features. However, no clear pattern is observed in sc-methylation data. This is likely because sc-methylation data is very noisy and over 20 percentage of the entries are missing in sc-methylation data. There is no clear pattern in sc-methylation data even if we group the cells by the provided cell type labels (Fig.11). Overall, our method *couple*CoC has better clustering results, likely due to the transfer of knowledge from clustering the scRNA-seq data to clustering sc-methylation data.

#### 3.2.3 Example 3: clustering pulmonary alveolar type 2, clara and ependymal cells of human and mouse scRNA-seq data

The last coupled real dataset includes human and mouse scRNA-seq data (Fran et al., 2019), and two cases are considered: (A) 99 clara cells and 14 ependymal cells in the mouse scRNA-seq dataset, and 113 clara cells and 58 ependymal cells in the human scRNA-seq dataset; (B) other than the cells in the first case, 179 mouse pulmonary alveolar type II cells were added, so there is one cell type in the target data (mouse) that is not present in the source data (human). We use the cell-type annotation (Angelidis et al., 2019) as a benchmark for evaluating the performance of the clustering methods. Fig.10 shows the optimal number of clusters are *N*_T_ = 2 and *N*_S_ = 2 for both cases, and for the target data in the case B, values of Calinkski-Harabasz index are close when the numbers of clusters are 2 or 3. Here we use the true number of cell clusters *N*_T_ = 3 instead of *N*_T_ = 2 in case B. The tuning parameters in *couple*CoC are set as λ_1_ = 1, λ_2_ = 0.2, *K* = 5 by grid search. Table 4 shows that in case A, SC3 performs well on source data, but it does not perform well on target data. *couple*CoC and Seurat perform better than the other methods for clustering the target data and the source data. In case B, *couple*CoC performs the best for clustering the target data. Since *N*_T_ > *N*_S_ in this case, we postprocessed the cluster labels to match the clusters across the two datasets, following the steps in Section 2.2. Table 5 shows that the cell types are matched by *couple*CoC. This example demonstrates that even when the cell types are not perfectly matched in the source data and the target data, *couple*CoC still works well and it improves the clustering of target data by effectively transferring knowledge from source data. Fig.4 shows the heatmap after clustering by *couple*CoC. Our proposed *couple*CoC clearly clusters similar cells and features. The datasets in example 3 are very sparse because they are generated on the drop-seq platform. Lastly, we also present the heatmap of clustering by *couple*CoC in case B with *N*_T_ = 2 in Fig.12, and it demonstrates that *couple*CoC can also give good clustering results.

**Figure 4:**
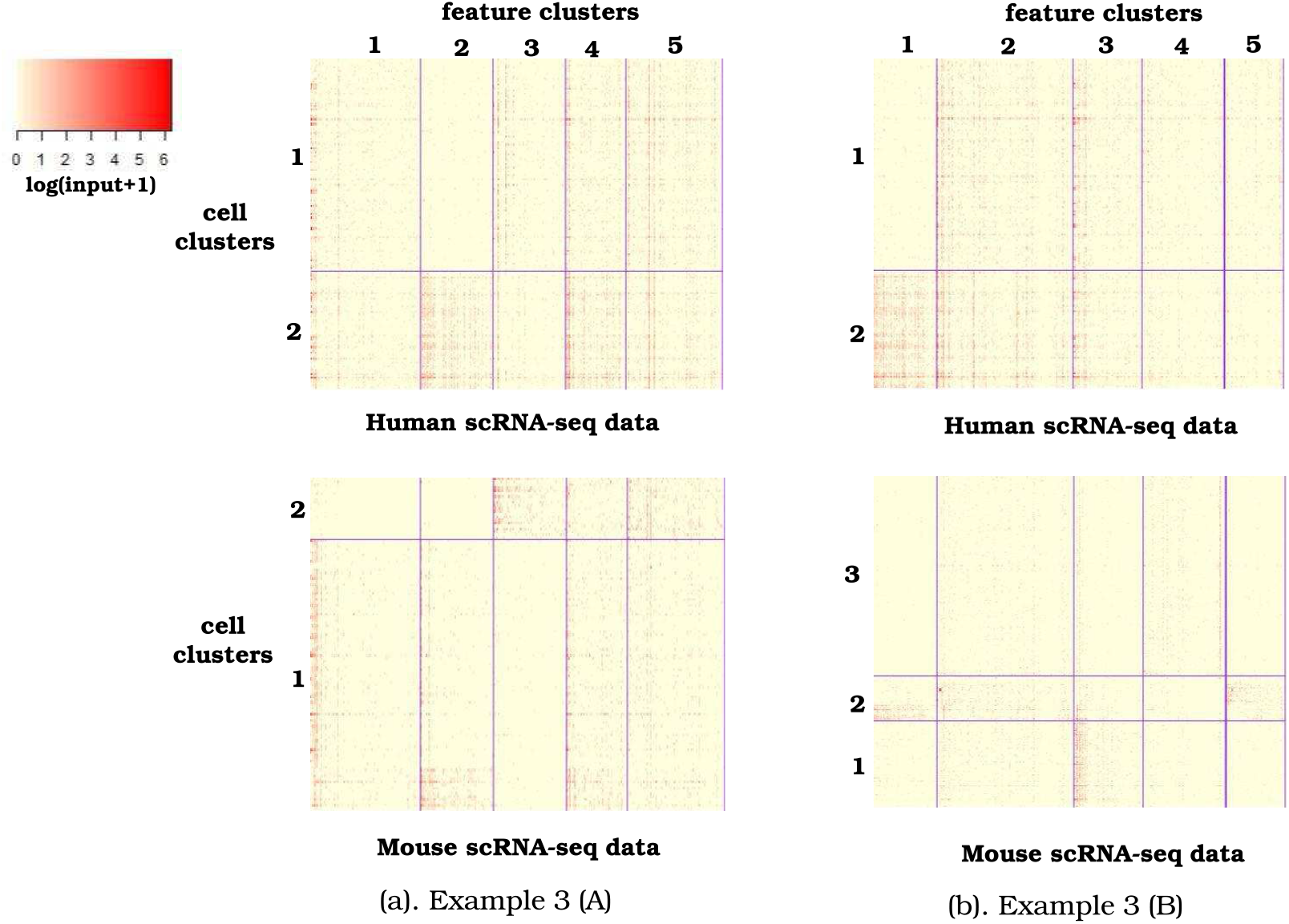
Heatmaps of the clustering results by *couple*CoC for example 3. The input stands for TPM. Case A: *N*_T_ = 2, *N*_S_ = 2. Case B: *N*_T_ = 3, *N*_S_ = 2.

**Table 4:** The results of clustering human and mouse scRNA-seq data in two cases. “T” and “S” represent target data and source data, respectively.

| Settings | Methods | Purity | RI | ARI | NMI |
| --- | --- | --- | --- | --- | --- |
| Example 3 (A)<br>($N_T = 2$)<br>($N_S = 2$) | *coupleCoC* (T) | 0.876 | 0.569 | -0.105 | 0.065 |
|  | *k*-means (T) | 0.823 | 0.706 | 0.053 | 0.006 |
|  | SC3 (T) | 0.796 | 0.673 | -0.083 | 0.029 |
|  | SIMLR (T) | 0.823 | 0.706 | 0.358 | 0.279 |
|  | SOUP (T) | 0.814 | 0.695 | 0.087 | 0.016 |
|  | Seurat (T) | 0.876 | 0.781 | 0.000 | 0.000 |
|  | *coupleCoC* (S) | 0.982 | 0.965 | 0.930 | 0.883 |
|  | *k*-means (S) | 0.778 | 0.652 | 0.268 | 0.190 |
|  | SC3 (S) | 0.988 | 0.977 | 0.953 | 0.908 |
|  | SIMLR (S) | 0.848 | 0.741 | 0.481 | 0.431 |
|  | SOUP (S) | 0.994 | 0.988 | 0.976 | 0.948 |
|  | Seurat (S) | 0.965 | 0.932 | 0.862 | 0.783 |
| Example 3 (B)<br>($N_T = 3$)<br>($N_S = 2$) | *coupleCoC* (T) | 0.914 | 0.895 | 0.790 | 0.672 |
|  | *k*-means (T) | 0.856 | 0.781 | 0.563 | 0.460 |
|  | SC3 (T) | 0.890 | 0.920 | 0.841 | 0.738 |
|  | SIMLR (T) | 0.801 | 0.821 | 0.643 | 0.573 |
|  | SOUP (T) | 0.877 | 0.831 | 0.664 | 0.524 |
|  | Seurat (T) | 0.887 | 0.848 | 0.697 | 0.508 |
|  | *coupleCoC* (S) | 0.982 | 0.965 | 0.930 | 0.883 |

**Table 5:** Matching the cell types by *couple*CoC in the case B of example 3. “Clu” is the abbreviation of “Cluster”, and “paII” stand for “pulmonary alveolar type II”.

| Example 3<br>(B) |  | *coupleCoC* |  |  | Purity | after postprocessing |  |  | Purity |
| --- | --- | --- | --- | --- | --- | --- | --- | --- | --- |
|  |  | Clu1 | Clu2 | Clu3 |  | Clu1 | Clu2 | Clu3 |  |
| mouse scRNA-seq data<br>(T, $n_T = 292$) | clara | 4 | 21 | 74 | 0.914 | 74 | 21 | 4 | 0.914 |
|  | ependymal | 0 | 14 | 0 |  | 0 | 14 | 0 |  |
|  | paII | 172 | 5 | 2 |  | 2 | 5 | 172 |  |
| human scRNA-seq data<br>(S, $n_S = 171$) | clara | 110 | 3 | | 0.982 | 110 | 3 | | 0.982 |
|  | ependymal | 0 | 58 |  |  | 0 | 58 |  |  |

## 4 Conclusion

Unsupervised methods, including dimension reduction and clustering, are essential to the analysis of single-cell genomic data as the cell types need to be inferred from the data. In addition, the matching of cell types in multimodal data cannot be disregarded, because it can provide rich fucntional information. In this work, we have developed an unsupervised transfer learning approach for the integrative analysis of multimodal single-cell genomic data, and we have demonstrated its performance on simulated data and real data analysis. Through the linked features across multimodal single-cell data, our proposed method can effectively transfer knowledge across multiple datasets. The degree of knowledge transfer is learnt adaptively from the datasets. Our proposed method is applicable in other areas including text mining.

## Appendices

## Supplementary Text

Similar to the simulation scheme designed by (Lin et al., 2019), we generate data in simulation study as follows:

1. Generate **w**^*acc*^ and **w**^*exp*^.

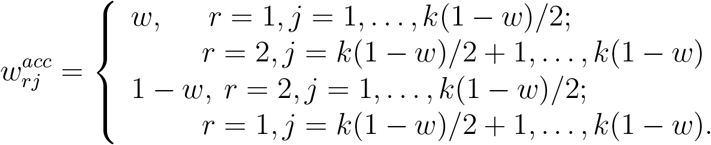

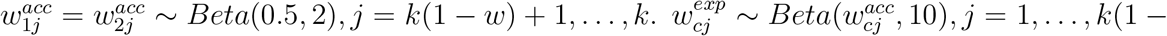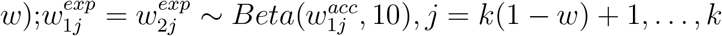.
2. Generate *z*^*acc*^ and *z*^*exp*^. The cluster labels are generated with equal probability 0.5.
3. Generate *u*^*acc*^ and 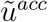. 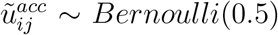, if 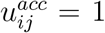, where 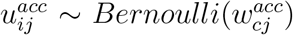 if 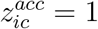, 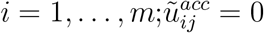 otherwise.
4. Generate *u*^*exp*^ and 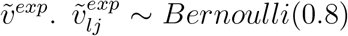 if 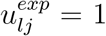, where 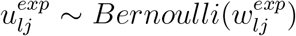 if 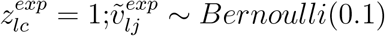 otherwise; *l* = 1, …, *n*.
5. Generate *C* and *G*. *C*_*ij*_ ~ *N*(0, 0.6^2^) if 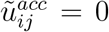 and *C*_*ij*_ ~ *N*(2, 0.6^2^) if 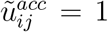; *G*_*lj*_ ~ *N*(0, *σ*^2^) if 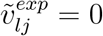 and *G*_*lj*_ ~ *N*(2, *σ*^2^) if 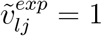.
6. Generate T and S. T_*lj*_ = 1 if G_*lj*_ > 0 and T_*lj*_ = 0 otherwise; S_*ij*_ = 1 if *C*_*ij*_ > 0 and S_*ij*_ = 0 otherwise.

More details on the notations and the simulation scheme is presented in Lin et al. (2019). In our simulation, the difference on the steps of data generation from that in Lin et al. (2019) is the addition of Step 6, which generates binary data. T_*lj*_ = 1 means gene *j* is expressed in cell *l*, and T_*lj*_ = 0 otherwise. S_*ij*_ = 1 means the promoter region for feature *j* is accessible in cell *i*, and S_*ij*_ = 0 otherwise.

## Supplementary Figures

**Figure 5:**
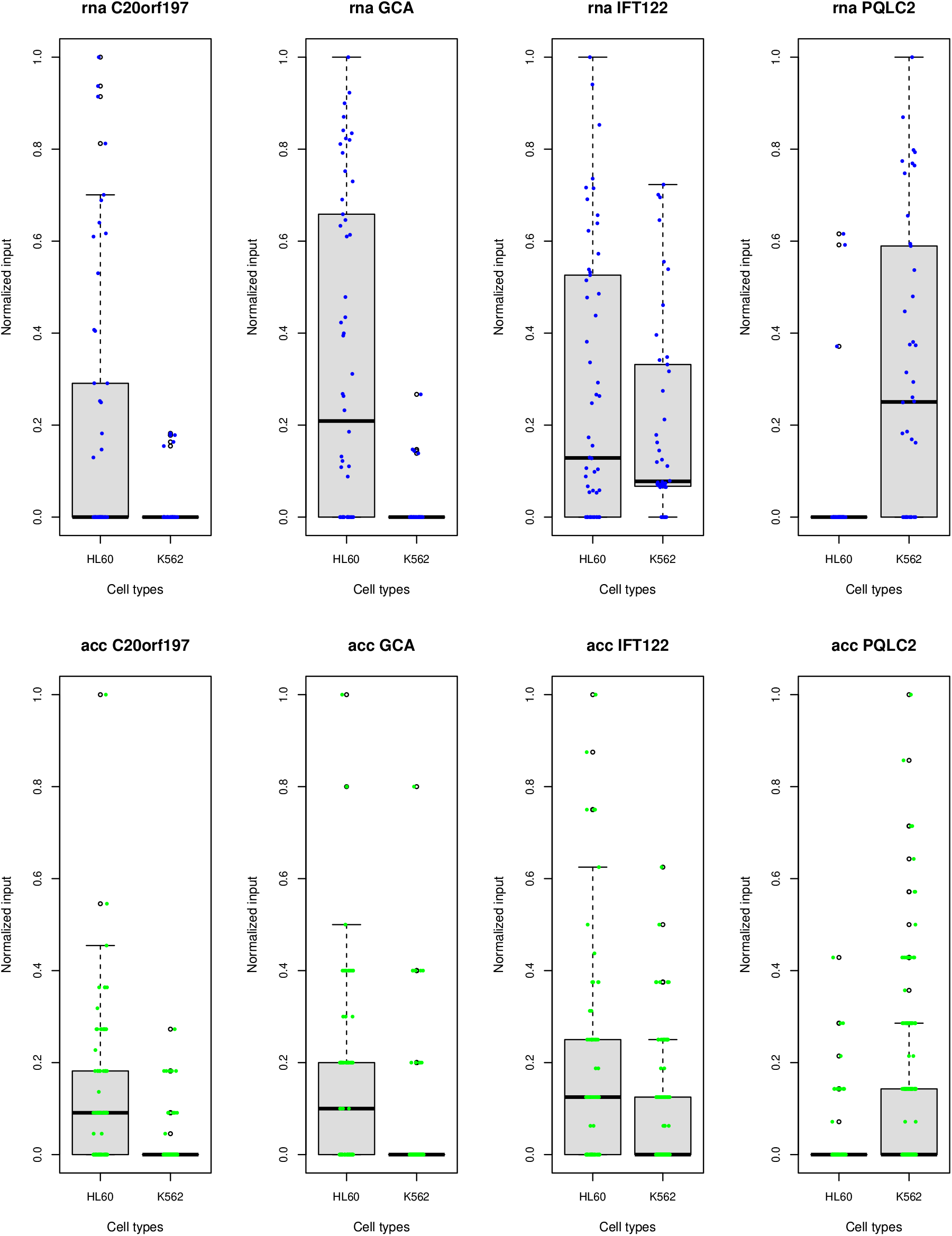
Mixture of jitter plots and boxplots to show the distribution of four features selected for demonstrating the positive relationship of features between scRNA-seq data and scATAC-seq data in example 1 (A).

**Figure 6:**
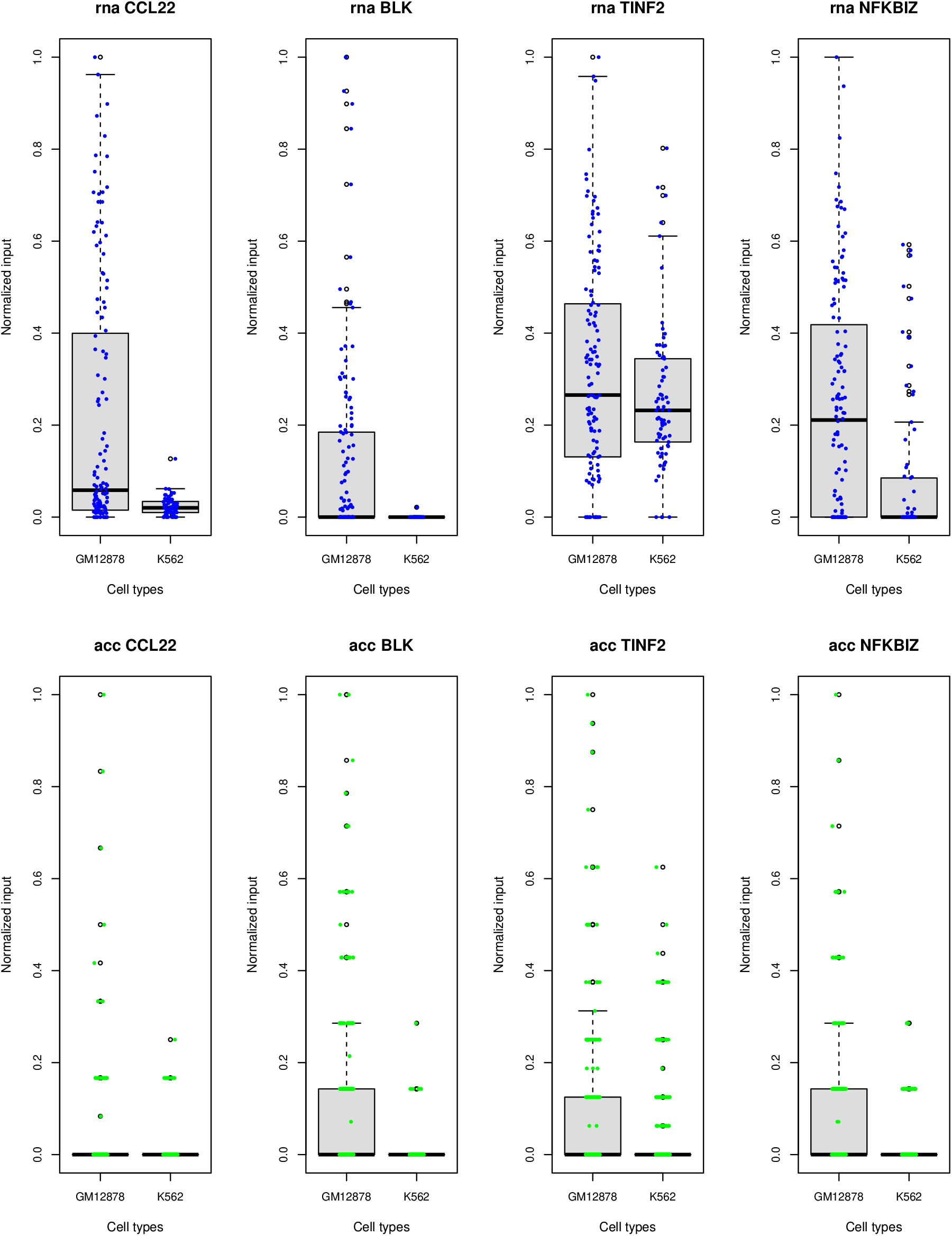
Mixture of jitter plots and boxplots to show the distribution of four features selected for demonstrating the positive relationship of features between scRNA-seq data and scATAC-seq data in example 1 (B).

**Figure 7:**
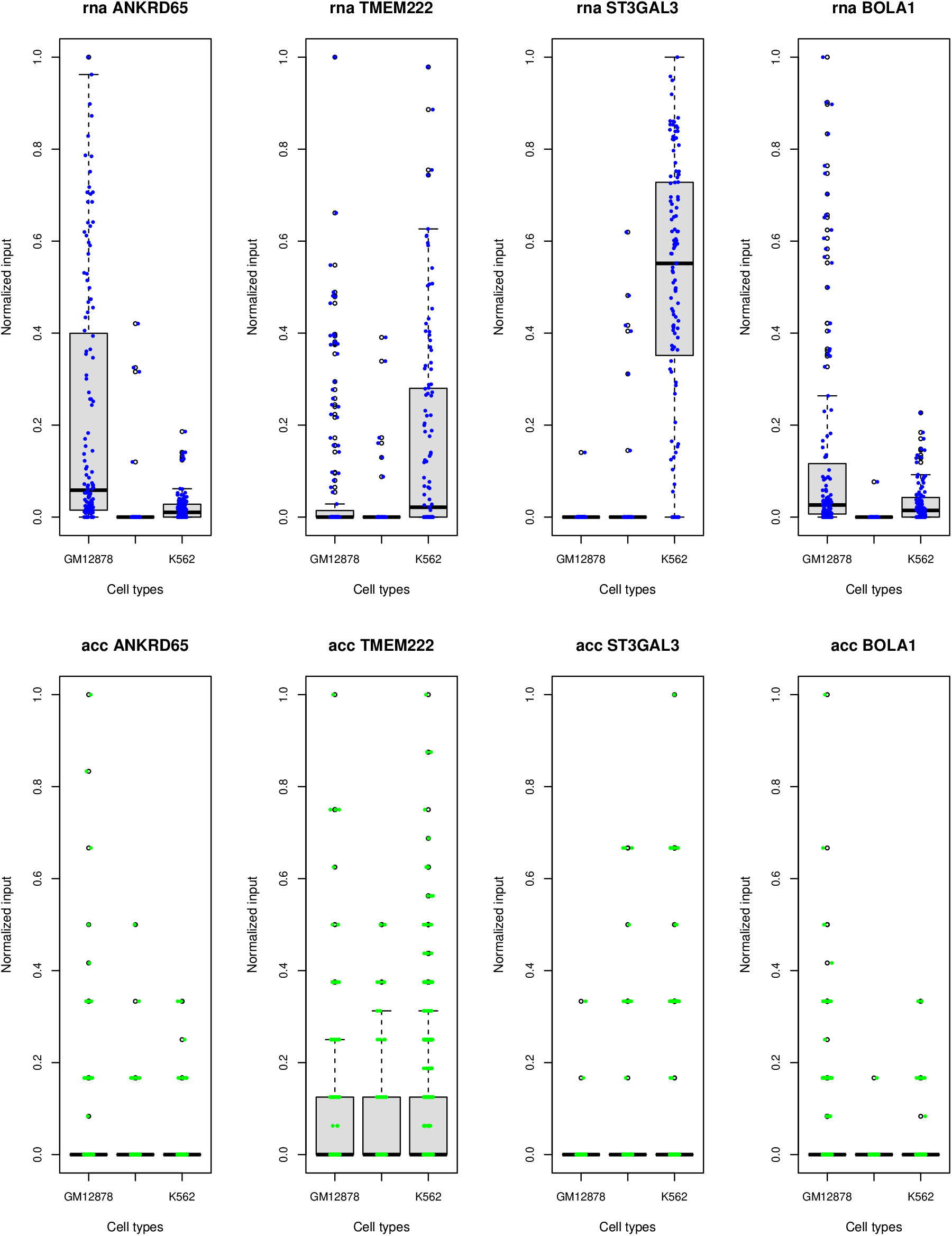
Mixture of jitter plots and boxplots to show the distribution of four features selected for demonstrating the positive relationship of features between scRNA-seq data and scATAC-seq data in example 1 (C).

**Figure 8:**
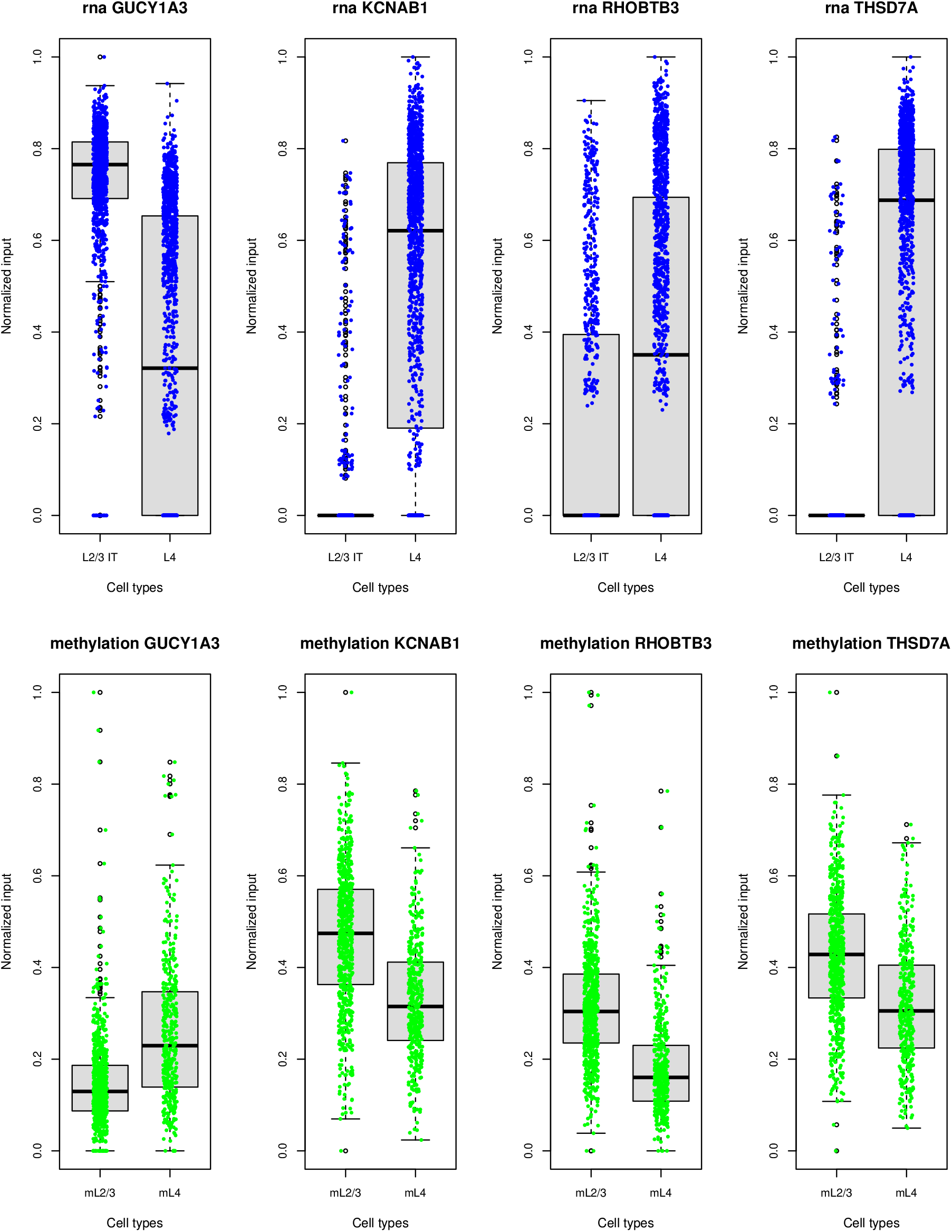
Mixture of jitter plots and boxplots to show the distribution of four features selected for demonstrating the negative relationship of mouse sc-methylation data and scRNA-seq data in example 2.

**Figure 9:**
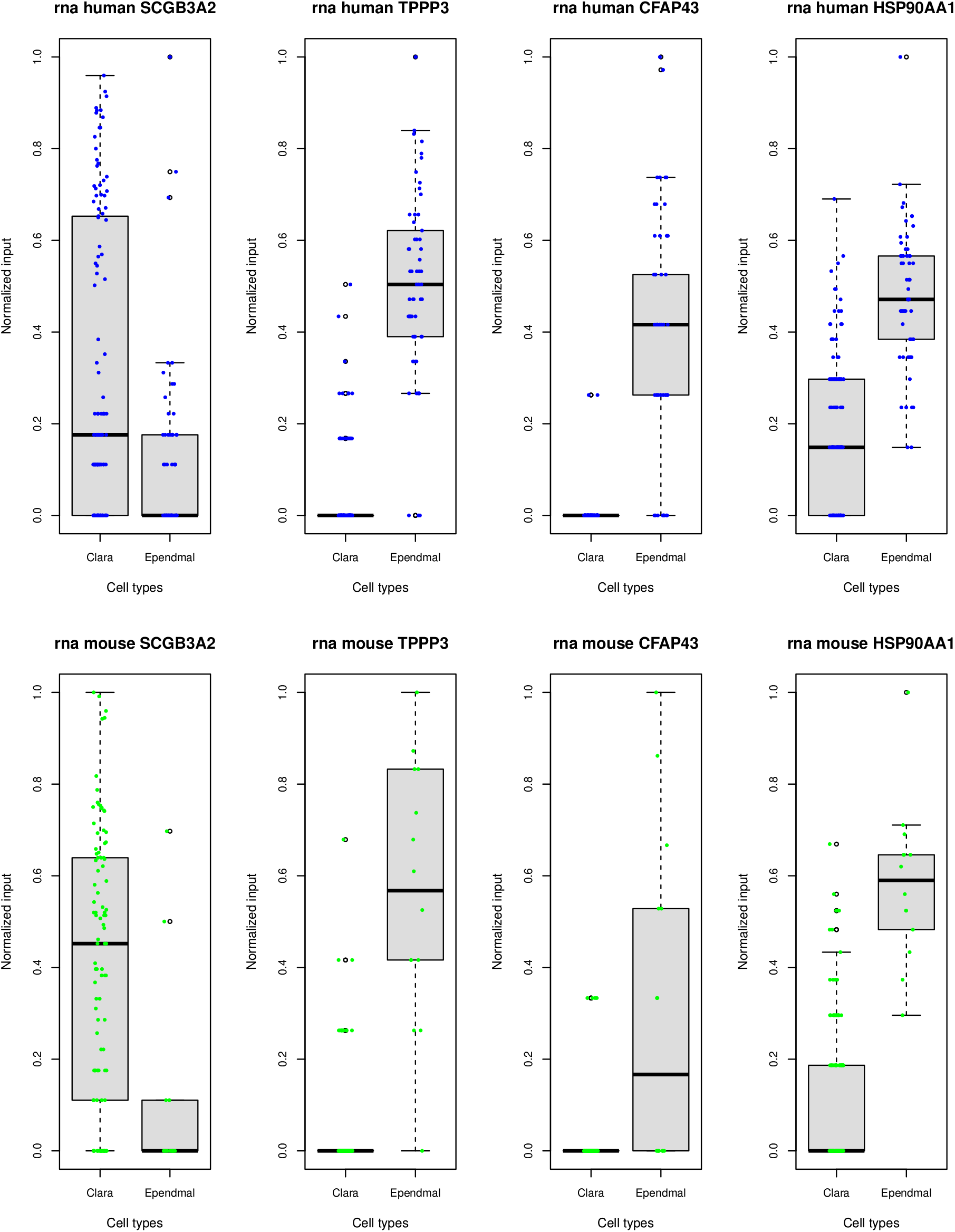
Mixture of jitter plots and boxplots to show the distribution of four features selected for demonstrating the positive relationship of homologs between human and mouse scRNA-seq data in example 3.

**Figure 10:**
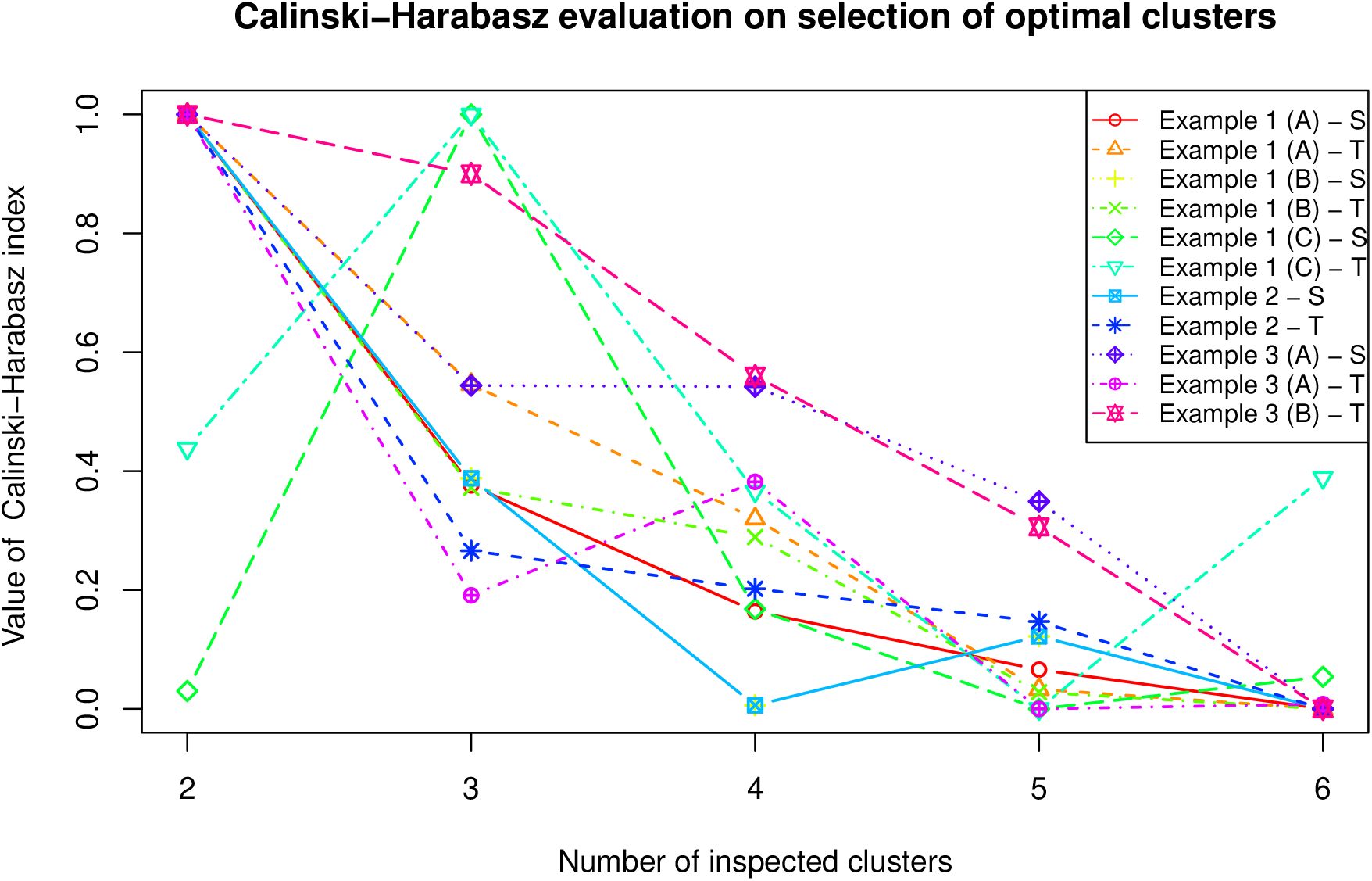
Calinski-Harabasz evaluation on selecting the optimal number of cell clusters for coupled datasets in real data examples. The value of Calinski-Harabasz index has been standardized via minimax normalization to ensure each value being bound to between 0 and 1 for each dataset in each example.

**Figure 11:**
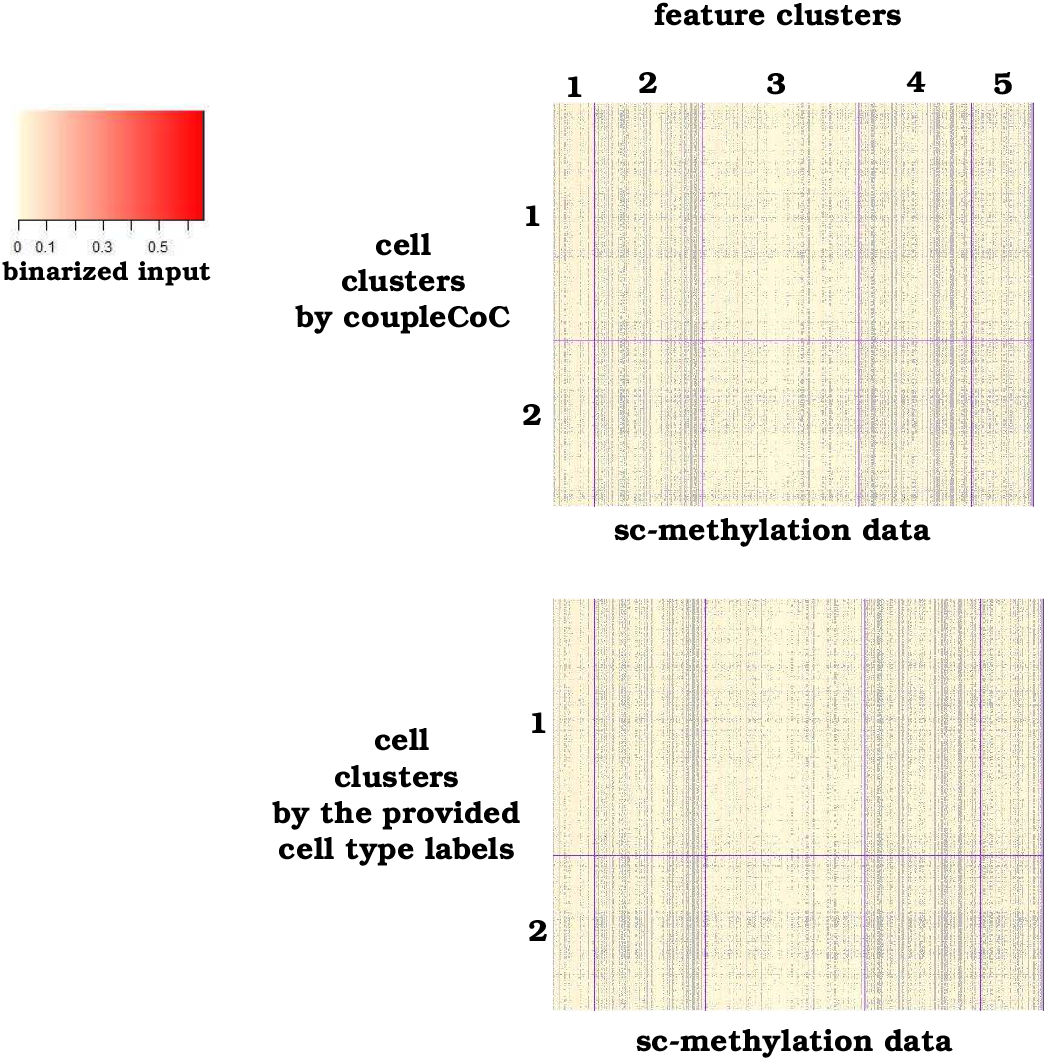
Heatmaps of clustering sc-methylation data by *couple*CoC and by the provided cell type labels for example 2.

**Figure 12:**
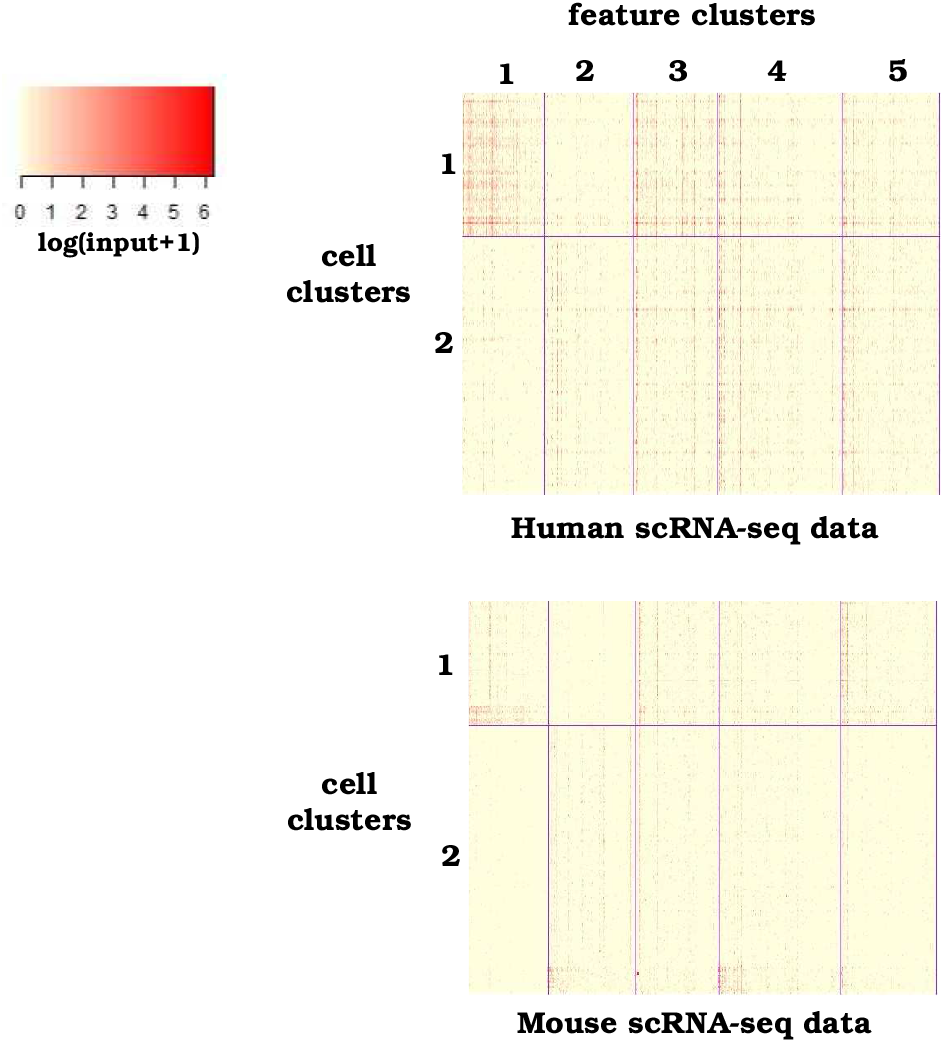
Heatmaps of the clustering results by *couple*CoC for case B of example 3, with the given number of cell clusters *N*_T_ = 2, *N*_S_ = 2.

